# Development of a novel hybrid alphavirus-Nipah virus pseudovirion for rapid quantification of neutralization antibodies

**DOI:** 10.1101/2025.02.17.638738

**Authors:** Brian Hetrick, Sravya Sowdamini Nakka, Dongyang Yu, Sarah Swineford, Nadia Storm, Anthony Griffiths, Yuntao Wu

**Author notes:** American Type Culture Collection (ATCC), Manassas, VA 20110, USA. Bond Life Sciences Center, Laboratory for Infectious Disease Research and Department of Molecular Biology, Microbiology and Immunology, University of Missouri; Columbia, MO 65211, USA.

## Abstract

Nipah virus (NiV) is an emergent paramyxovirus that causes serious disease in humans and animals. Accurate quantification of neutralizing antibodies (nAB) against NiV is essential for vaccine development. The current standard, the plaque reduction neutralization test (PRNT), requires authentic infectious NiV and BSL-4 containment, taking 4-7 days to complete. In this study, we developed a hybrid alphavirus-Nipah virus pseudovirion (Ha-NiV) composed of a non-replicating Nipah virus virus-like particle (VLP) that encapsulates an RNA genome from a fast-expressing SFV (Semliki Forest Virus) alphaviral vector. Ha-NiV can infect NiV target cells but is replication incompetent, enabling rapid quantification of nAB (6-18 hours) in BSL-2 conditions. We validated Ha-NiV using sera from Nipah virus-infected African green monkeys and established a good correlation (R^2^ = 0.86) with PRNT, demonstrating comparable specificity and sensitivity. These results demonstrate that the Ha-NiV assay can serve as a novel platform for convenient and rapid quantification of nAB in BSL-2 settings.

## INTRODUCTION

Nipah Virus (NiV) infection was first reported in Malaysia in 1998. The outbreak in pig farms caused 229 human cases of febrile encephalitis with 111 deaths (48% fatality). The viral pathogen was subsequently identified in 1999 and named after the village of Nipah, where the first human infection was reported ^1–3^. NiV is a highly pathogenic virus that belongs to the Paramyxoviridae family, specifically within the genus Henipavirus ^4^. The reservoir species for NiV are fruit bats and humans typically become infected via contact with infected animals (such as pigs) or contaminated food (raw date palm sap) ^5^. Following the identification of NiV, numerous outbreaks have occurred in Southeast Asia. For example, a 2001 outbreak in West Bengal, India resulted in 66 cases and 45 deaths. Recently, in 2023, two outbreaks in Bangladesh and India also resulted in 11 cases and two deaths in one incidence, and at least five cases and two deaths in the other ^6^. Because of the high mortality rate (40-75%), NiV was classified by CDC as a Category C agent. Currently, there is no effective treatment or approved vaccines for NiV.

NiV is an enveloped virus with a large 18 kb negative-sense, single-stranded RNA genome that contains six structural proteins: glycoprotein (G), attachment protein (F), nucleoprotein (N), matrix protein (M), phosphoprotein (P), and large protein (L) ^7^. G and F are essential for viral entry, with G facilitating binding to cell receptors (Ephrin-B2) ^8^ and F mediating the fusion of the viral envelope with the host cell membrane ^9^. N encapsulates the viral RNA genome ^10^, while M provides structural stability for virion assembly ^11^. P interacts with L and N and plays a role in viral RNA synthesis and modulating host responses ^12–15^, and L serves as the RNA polymerase responsible for replicating and transcribing viral mRNA ^12^.

Vaccines for NiV are currently in various stages of development, with several promising candidates emerging. Some of these vaccines employ live attenuated recombinant VSV (vesicular stomatitis virus) to express G and F as immunogens ^16,17^, while others utilize mRNA or subunit vaccines that use the G and F proteins as the antigens ^18–20^. Additionally, viral vectors such as recombinant VSV ^16^ and modified chimpanzee adenoviral vector (ChAdOX1 NipahB) are used to express the G and F proteins as immunogens ^21^. In particular, ChAdOx1 NipahB has demonstrated robust immune responses and protection against NiV in preclinical animal models, and has commenced phase I clinical trials in January 2024 ^21,22^. The mRNA vaccine candidate mRNA-1215 is also in phase I clinical trial ^23^.

Evaluation of vaccine efficacy in animal and clinical trials is critical for vaccine development. Currently, the plaque reduction neutralization test (PRNT) is the standard method for evaluation of neutralizing responses induced by NiV vaccines ^24^. However, the assay requires the use of authentic infectious Nipah virus and BSL-4 containment, and also takes 4-7 days to complete, which limits large-scale and timely evaluations of neutralizing antibodies. Pseudovirus systems based on lentivirus (HIV, human immunodeficiency virus) ^25,26^, retrovirus (MuLV, moloney murine leukemia virus) ^27^, or VSV ^28–30^ have been developed to allow the assays to be performed using BSL-2 conditions in 1-2 days. These systems have provided very valuable tools and enabled the evaluation of vaccine candidates both *in vitro* and in animal models using non-BSL-4 conditions ^26^. Nevertheless, both VSV- and lentiviral-based pseudoviral particles contain only the NiV G and F proteins, and sometimes in truncated forms ^25^. The replacement of authentic NiV matrix protein with the structural proteins of VSV or lentivirus in these particles may affect viral assembly and virion properties in response to antibody neutralization ^11,31^. For example, when compared with PRNT, the VSV-NiV-F/G-based assay has been shown to be less effective in detecting low levels of antibodies (e.g. 47% sensitivity) ^29^. An important issue for VSV-based pseudoviruses is the presence of residual VSV, which can result in false-positive results ^29,32^. In addition, attenuated, G/F-expressing recombinant VSV is actively being developed as NiV vaccines ^16,17^, which also induce antibodies against VSV’s virion proteins. This may significantly impact the use of VSV-based pseudoviruses in anti-NiV neutralizing antibody assays. To overcome the limitations of the current NiV pseudovirus systems, here we describe the development of a new hybrid alphavirus-NiV pseudoviral particle (Ha-NiV) for rapid quantification of neutralization antibodies. Ha-NiV is a non-replicating virus-like particle composed of authentic NiV structural proteins (G, F, M, and N) with no structural proteins from other viruses. Ha-NiV also contains a self-amplifying RNA genome derived from an alphavirus-based vector ^33,34^ that can rapidly and robustly express reporter genes within 6 hours after viral entry ^34^. In this study, we also validated Ha-NiV with authentic virus PRNT and demonstrate that Ha-NiV can be used as a robust platform for rapid quantification of neutralization antibodies.

## RESULTS

To establish a rapid cell-based Nipah virus infection assay for screening and quantifying neutralizing antibodies and antiviral drugs in BSL-2 conditions, we developed a new hybrid alphavirus-Nipah pseudoviral particle, in which an alphavirus-based RNA genome is encapsulated for rapid expression of reporter genes in target cells (**Figure 1A**). The genomic RNA consists of the 5’ untranslated region and open-reading frames coding for the nonstructural proteins (nsp) 1-4 from Semliki Forest virus (SFV) ^33,35^. The nsp1-4 protein complexes mediate the self-amplification of the RNA genome in cells ^36^. The RNA genome also contains viral subgenomic RNA promoters for the expression of reporter genes such as *luciferase* (*Luc*) and *green fluorescent protein* (*GFP*). The downstream of the report genes includes the 3’ untranslated region of SFV and a poly(A) tail that are used to stabilize RNA.

**Figure 1.**
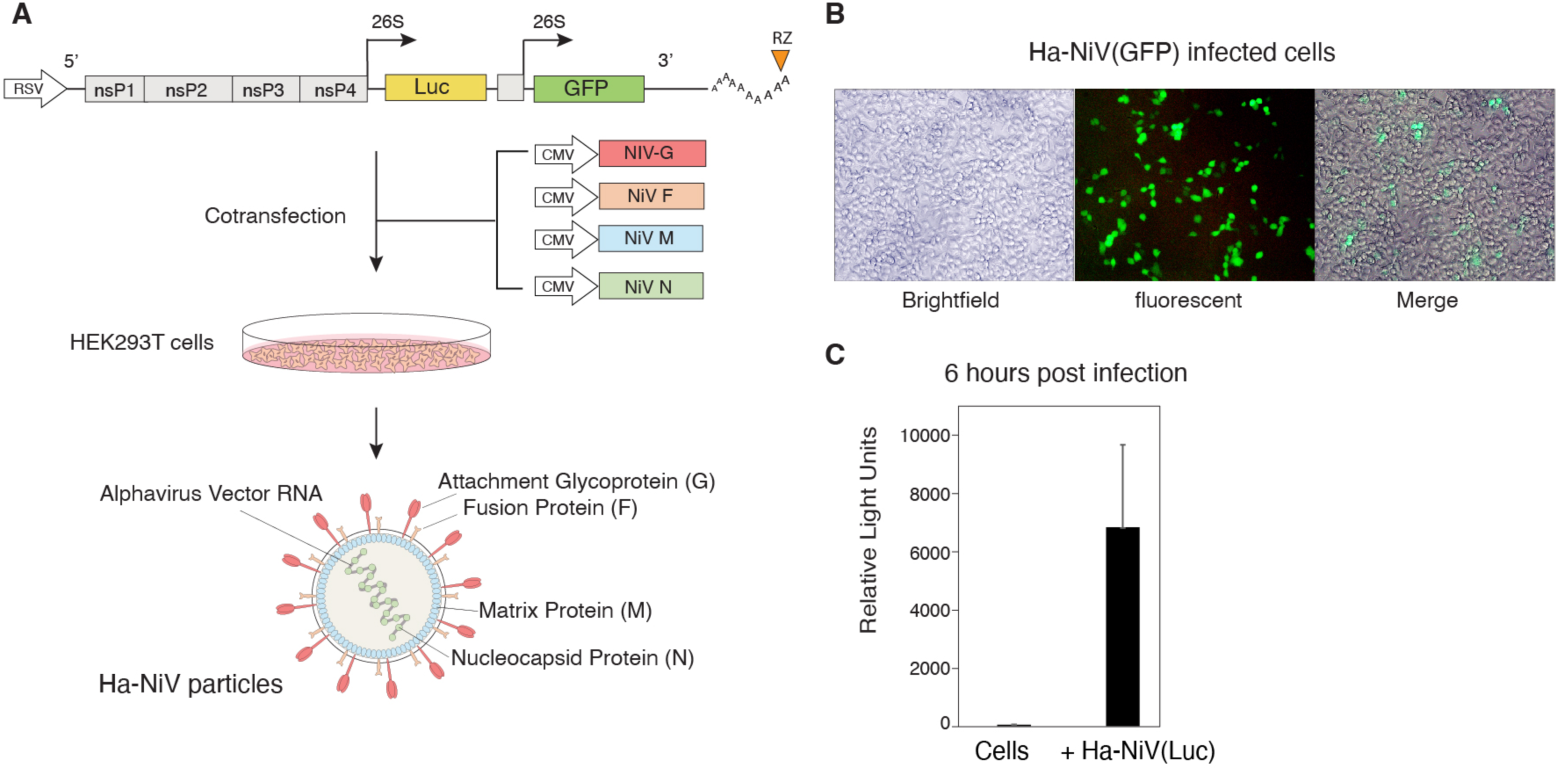
Design, assembly, and infection of Ha-NiV particles. **(A)** Illustration of the design and assembly of Ha-NiV vector and particles. The alphavirus-based genomic vector contains a RSV promoter that transcribes viral RNA genome to be packaged into Ha-NiV particles. Shown are the 5’ untranslated region followed by open-reading frames coding for nonstructural proteins (nsp) 1-4 from Semliki Forest virus (SFV), viral subgenomic promoters for Luc and GFP reporter expression, the 3’ untranslated region, and a poly(A) tail that is self-cleaved by the hepatitis delta virus ribozyme (RZ). To assemble viral particles, HEK293T cells were co-transfected with the Ha-Niv vector and the vectors expressing the 4 structural proteins of NiV (G, F, M, and N). Ha-NiV particles in the supernatant were harvested at 48 hours. (**B**) Infection of target cells with Ha-NiV(GFP) particles. HEK293T cells were infected with Ha-NiV (GFP) particles. GFP expression was observed 24 hours post infection. (**C**) Rapid reporter expression following Ha-NiV infection. Vero cells were infected with Ha-NiV(Luc) for 6 hours, washed, and then lysed and analyzed for Luc expression. The addition of virus to cells was defined as time “0”. Infection and luciferase assays were performed in 3 times, and the mean and standard deviation (SD) are shown.

To assemble viral particles, we used the Ha-NiV DNA vector to express the genomic RNA, and cotransfected the vector with vectors expressing the structural proteins of Nipah virus, which include the surface glycoproteins for attachment (G) and fusion (F), the matrix (M), and the nucleocapsid (N) (**Figure 1A**). Virion particles were harvested at 48 hours post cotransfection, and tested for virion infectivity by quantifying reporter gene expression in target cells. As shown in **Figure 1B**, infection of HEK293T cells with the Ha-NiV(GFP) particles led to robust GFP expression at 24 hours post infection, demonstrating the capacity of Ha-NiV(GFP) to enter and express genes in target cells.

A major advantage for utilizing alphavirus-based RNA genome for Ha-NiV is the capacity of the alphavirus-based genome for self-amplification, which can lead to extremely fast and high levels of gene expression; it has been shown that during alphavirus infection, gene expression from the subgenomic RNA promoters occurs within hours, and levels of viral plus-RNAs can reach 200,000 copies in a single cell ^33,34^. We assembled a Ha-NiV(Luc) particle, and followed the particle infection, and observed that the Luc reporter expression can be rapidly detected in 6 hours (**Figure 2C**). This rapid reporter expression permitted us to utilize Ha-NiV for rapid quantification of neutralization antibodies.

**Figure 2.**
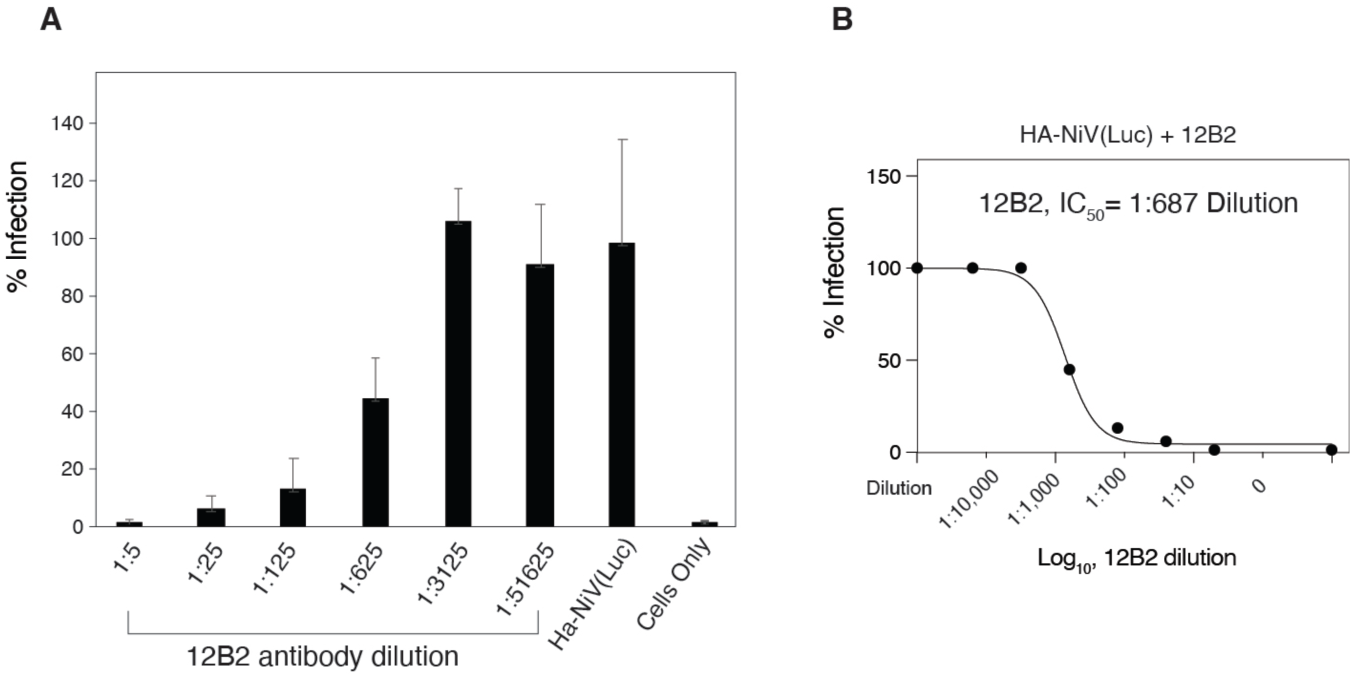
Validation of Ha-NiV(Luc) particles for rapid quantification of neutralizing antibodies. **(A)** Quantification of neutralizing antibodies with Ha-NiV(Luc) particles. Shown are the concentration-dependent inhibition of Ha-NiV(Luc) by a recombinant monoclonal antibody (12B2) to NiV F glycoprotein. 12B2 was serially diluted and incubated with Ha-NiV(Luc) particles for 1 hour at room temperature. The Ha-NiV(Luc)-antibody complex was used to infect HEK293T cells. Neutralization activities were quantified by luciferase assay at 18 hours post addition of virus to cells. (**B**) The IC_50_ was calculated using the relative percentage of infection with or without 12B2 neutralization.

To validate Ha-NiV for rapid quantification of neutralizing antibodies, we tested a commercial anti-NiV F glycoprotein antibody (12B2), which was serially diluted and pre-incubated with Ha-NiV(Luc). The antibody-virus complex or a control non-neutralized Ha-HiV were used to infect cells for 18 hours for Luc expression. As shown in **Figure 2A**, we observed antibody concentration-dependent inhibition of Ha-Niv(Luc), and the IC_50_ (half maximal inhibitory concentration) was determined to be at 1:687 dilution (**Figure 2B**).

Based on the results described above, we performed additional validation of Ha-NiV neutralizing assays using sera from Nipah virus-infected African green monkeys (total n =16, infected with various virus doses using different inoculation routes) that were part of other studies (unpublished). Serum samples collected at days 8, 9, 10, 12, 36, or 37 were quantified for the titers of neutralizing antibodies using Ha-NiV(Luc). In the nAB assays, serum samples were serially diluted and pre-incubated with Ha-NiV(Luc). The complex was then used for infecting HEK293T cells. Luc reporter expression was quantified at 18 hours post infection. We also included a standard nAB as the positive control and uninfected monkey sera as the negative control. As shown in **Figure 3**, the control standard antibody demonstrated antibody-concentration dependent inhibition of Ha-HiV(Luc) with an IC_50_ of 1:74,200 dilution, whereas the uninfected serum control did not show dosage-dependent inhibition. For the serum samples, the IC_50_ values were quantified at various levels, from 1:1.75×10^2^ to 1:3.15 x 10^9^. Several serum samples from monkeys infected with low levels of Nipah virus (1-100 PFU) (AGM-N05, AGM-N07, AGM-N08, AGM-N12, AGM-N13, AGM-N14, AGM-N15, AGM-N16) showed no antibody dosage-dependent inhibition of Ha-NiV(Luc), similar to the uninfected serum control.

**Figure 3.**
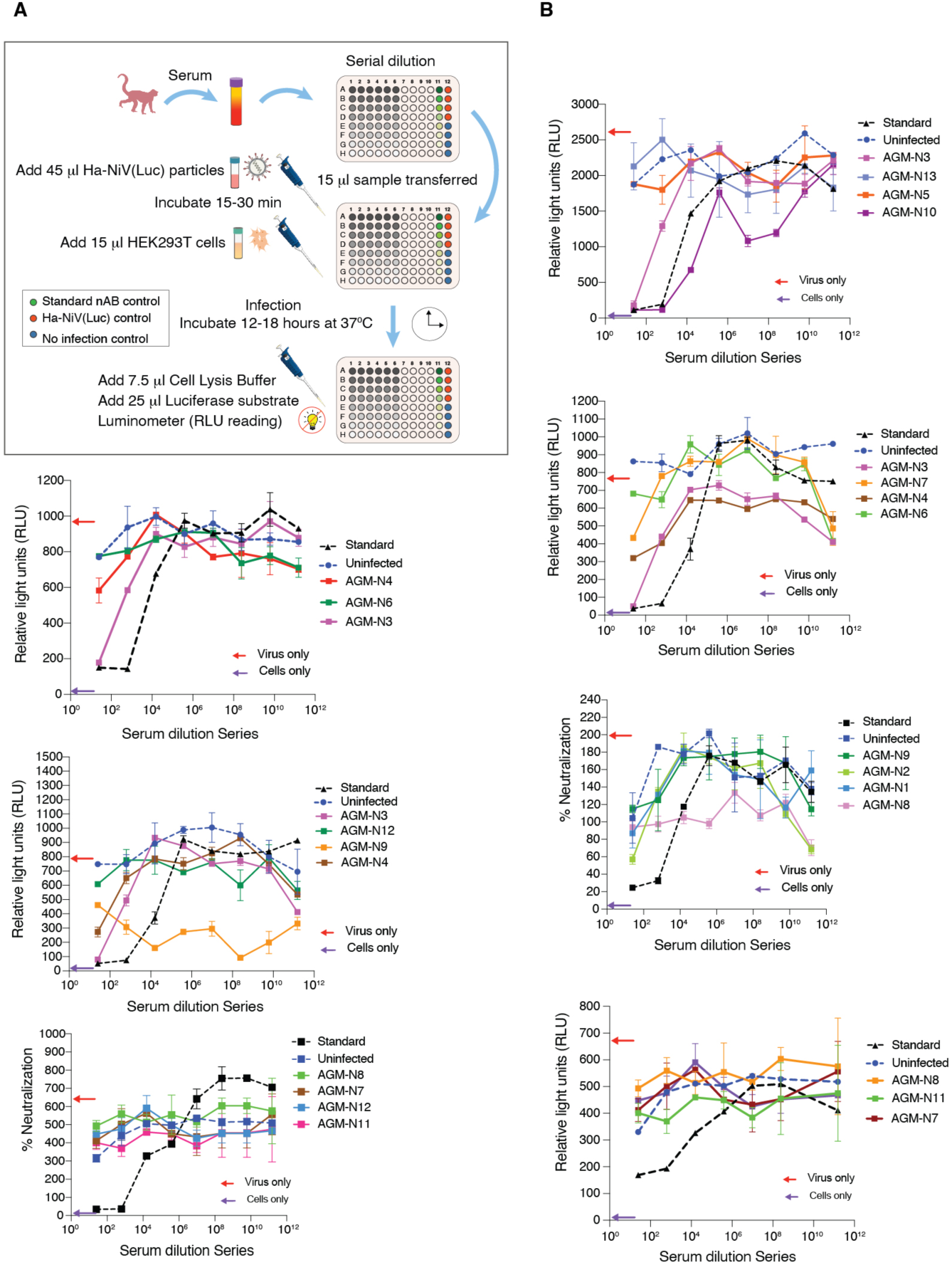
Validation of Ha-NiV(Luc) for rapid quantification of anti-sera from NiV-infected African green monkeys. (**A**) Illustration of the procedure for rapid quantification of anti-serum with Ha-NiV(Luc). (**B**) Quantification of neutralization activities of anti-sera with Ha-NiV(Luc) particles. Shown is the anti-serum concentration-dependent inhibition of Ha-NiV(Luc) by a standard neutralizing antibody (Standard) or the individual anti-sera from 16 NiV-infected animals. Neutralization activities were quantified by luciferase assay at 18 hours post addition of the antibody-virus complex to cells. Control serum was from uninfected animal sera (uninfected).

For comparison, we also performed an independent quantification of these serum samples using PRNT (**Figure 4**). We observed good correlations between PRNT and the Na-NiV(Luc) assay in specificity and sensitivity (F**ig. 5A**). When the IC_50_ titers were plotted, comparison of the titers for the PRNT and Ha-NiV(Luc) assay gave a correlation coefficient of 0.86 (**Figure 5B**). These results demonstrated that the Ha-NiV(Luc)-based nAB assay can be used for accurate quantification of neutralizing antibodies, similarly as PRNT, but with a much faster speed (6-18 hours versus 5-7 days) and without the need for BSL-4 containment.

**Figure 4.**
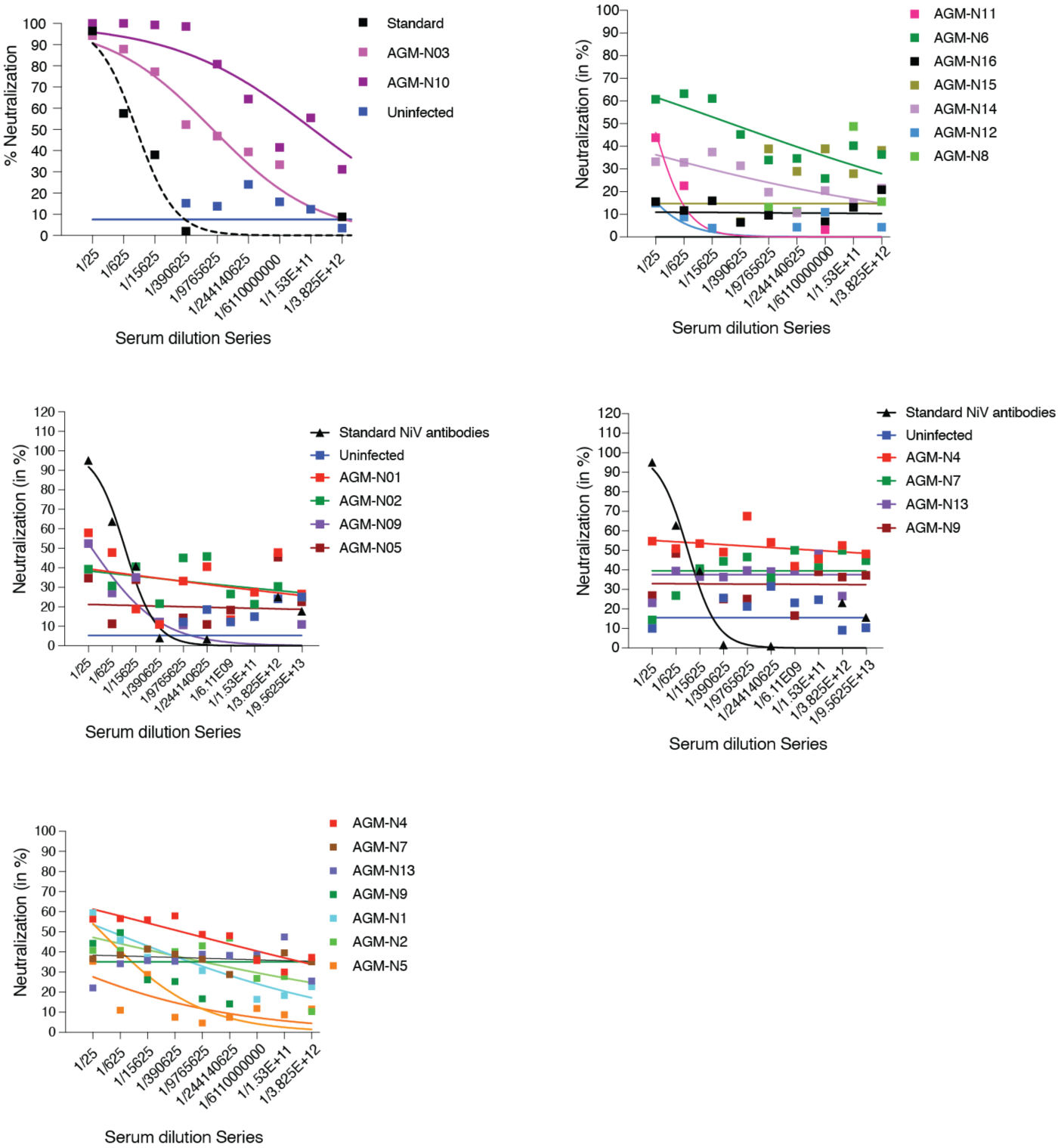
Quantification of anti-sera from NiV-infected African green monkeys using PRNT. For comparison with Ha-NiV nAB assay, the same anti-sera from 16 NiV-infected animals (Fig. 3B) were quantified for neutralization activities using the plaque reduction neutralization test (PRNT). Shown is the concentration-dependent reduction of NiV plaques by individual anti-sera or a standard neutralizing antibody (Standard). Uninfected animal sera (uninfected) were also used for negative control.

**Figure 5.**
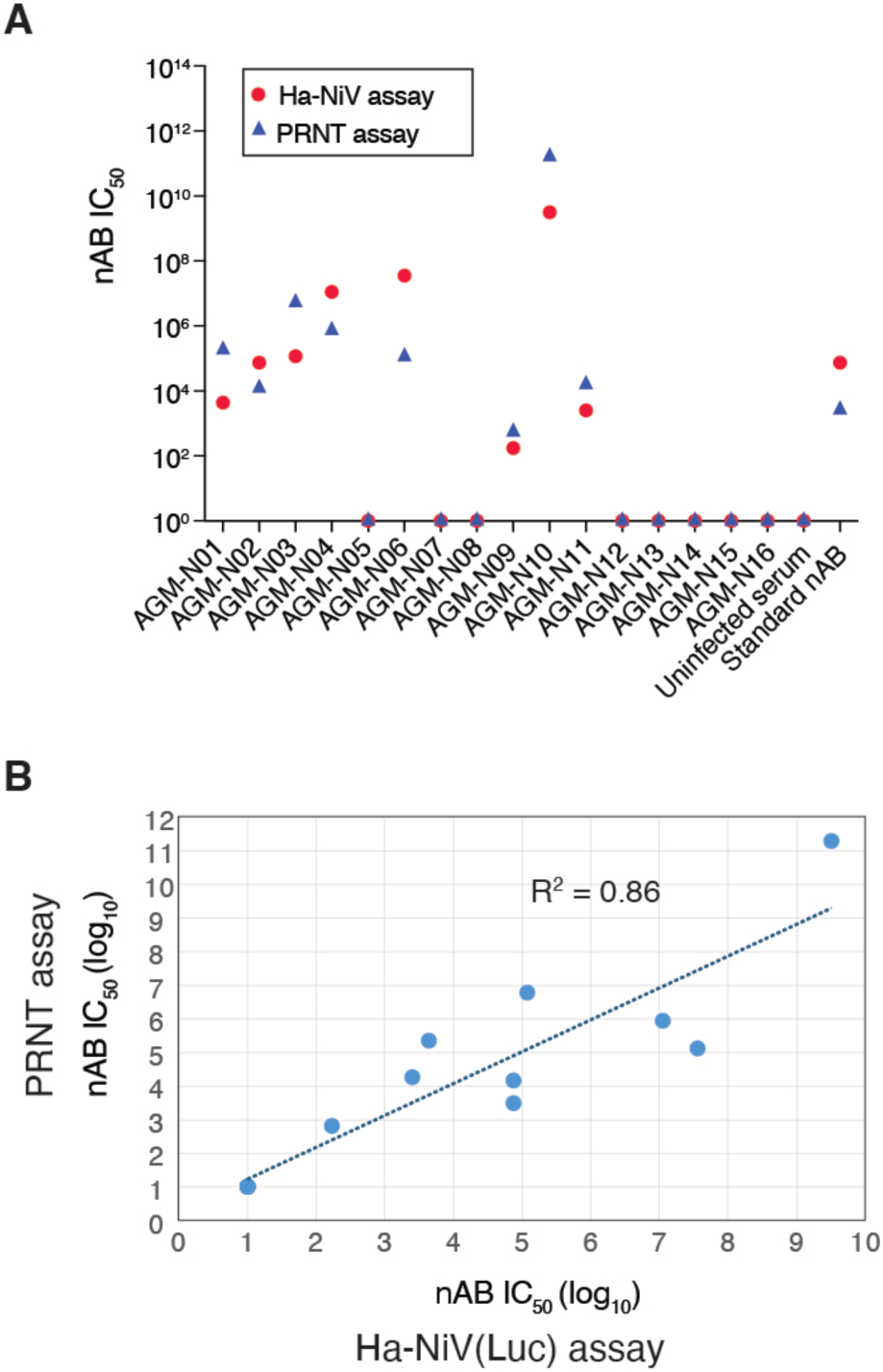
Correlation of serum neutralization activities quantified with Ha-NiV(Luc) and PRNT. **(A)** The neutralization activities of the anti-sera from 16 NiV-infected animals were quantified with the Ha-NiV(Luc) assay (Figure 3) or PRNT (Figure 4), and the IC_50_ values were calculated and plotted. (**B**) The IC_50_ values from (**A**) were plotted for the correlation of IC_50_ between the Ha-NiV(Luc) assay and PRNT.

## DISCUSSION

In this article, we describe the development and validation of a novel hybrid NiV pseudovirus system, the Ha-NiV particle. Structurally a virus-like particle (VLP), Ha-NiV retains the pseudovirus capability to enter target cells and express reporter genes. It incorporates a reporter genome from the robust alphavirus expression system, aligning with previous studies on alphaviral vectors. Alphaviral vectors are known for their high efficiency and rapid gene expression. For instance, Hahn *et al.* have demonstrated that these vectors produce high-titer particles (10⁸–10⁹ PFU/ml) in transfected cells and generate over 10⁶ CAT (chloramphenicol acetyltransferase) reporter molecules within 7 hours of infection ^37^. Similarly, Xiong et al. reported expression of 10⁸ CAT molecules per cell within 16-20 hours ^38^. Other studies have also shown that gene expression from subgenomic RNA promoters occurs within hours, with viral plus-RNA levels reaching up to 200,000 copies per cell ^33,34^.

We demonstrated that Ha-NiV can be effectively used for rapid screening and quantification of neutralizing antibodies. A direct comparison between the Ha-NiV assay and PRNT revealed a strong correlation (R^2^ = 0.86) for anti-serum quantification, validating Ha-NiV as a suitable surrogate for authentic infectious NiV in neutralizing antibody assays (**Fig. 5B**). However, PRNT, which uses infectious NiV, generally exhibits higher sensitivity than pseudovirus-based assays like the VSV-NiV-F/G assay ^29^. This discrepancy may stem from the presence of contaminating VSV particles in VSV-NiV-F/G preparations ^32^. These particles, containing largely VSV structural proteins, may not be neutralized by anti-NiV antibodies, potentially reducing assay sensitivity. In contrast, Ha-NiV offers a superior virion structure, forming native NiV virus-like particles composed exclusively of authentic NiV structural proteins (G, F, M, and N) without contamination from other viral proteins. Consequently, the Ha-NiV-based assay demonstrated comparable specificity and sensitivity to PRNT and exhibited a stronger correlation (R^2^ = 0.86) compared to the VSV-NiV-F/G-based assay (R^2^ = 0.51–0.68) ^29^.

The Nipah virus has been designated by the World Health Organization (WHO) as a critical priority pathogen, emphasizing the urgent need for accelerated research and development of new diagnostics, treatments, and vaccines ^39^. This classification arises from the virus’s alarming mortality rate and its potential to trigger large-scale outbreaks. The introduction of Ha-NiV represents a novel pseudovirus platform that not only accelerates the development of diagnostics and vaccines but also facilitates the identification of antiviral compounds, especially those aimed at disrupting viral entry.

## LIMITATIONS OF STUDY

The present method does have some limitations. For one, the Ha-NiV(Luc)/PRNT correlation study was only performed using monkey sera. Future studies may need to use human anti-sera for correlation studies. In addition, the expression of the Nipah virus structural proteins and their encapsulation of the alphaviral RNA genome may need to be further optimized.

## ACKNOWLEDGMENTS

We thank the Coalition for Epidemic Preparedness Innovations (CEPI) for funding support of the animal studies that provided the material used in this work. Authors would like to thank and acknowledge extensive animal work carried out at the NEIDL’s Animal Research Services (ARC), Boston University. This work was supported by Virongy Biosciences Inc. The PRNT correlation studies were supported by startup funds to A.G., Boston University.

## AUTHOR CONTRIBUTIONS

Research studies were designed by B.H., S.S.N., N.S., A.G., and Y. W.. Experiments were performed by B.H., S.S.N., D.Y., S.S. Manuscript was written by Y.W.

## METHODS

### Animals

Non-human primate work was performed in high containment laboratories at the NEIDL, Boston University (BU) and approved by BU’s Institutional Animal Use and Care Committee (IACUC) and conducted in accordance with Good Laboratories Practice (GLP) non-clinical laboratory studies. African Green Monkeys (AGMs) challenged with Nipah virus Bangladesh (NiVb) strain (200401066 Bangladesh, BEI resources, Manassas, VA, USA). Animals were exposed to varying NiVb doses (High, HD: 5.88E+04 PFU /ml; Medium-high, MD: 1.86E+04 PFU/ml; low, LD: 5.88E+02 PFU/ml) via intranasal (IN), intratracheal (IT) and both IN-IT routes of transmission. Animals were monitored for the disease progression and were euthanized when moribund while surviving animals were euthanized on the end of in-life phase set on day 36. Pooled serum from uninfected AGMs (n=2) was used as negative control.

### Virus and viral particle assembly

The NiV protein expression vectors (pCMV-NiV-G, pCMV-NiV-F, pCMV-NiV-M, pCMV-NiV-N) and the alphaviral vectors were provided by Virongy Biosciences Inc. Ha-NiV particles were assembled by cotransfection of HEK293T cells with NiV structural protein expression vectors (G, F, M, N) and the alphaviral Luc vector or alphaviral GFP vector. Particles were harvested at 48 hours post cotransfection, filtered through a 0.45 μm filter.

### Viral infectivity assay

Ha-NiV particles were used to infect HEK293T cells (ATCC). Briefly, cells were seeded in 12-well plates (2×10^5^ cells per well), and infected for 1-2 hours at 37°C, washed, cultured in fresh medium for 48 hours, and then observed for GFP expression, or lysed in luciferase lysis and assay buffer (Virongy) for luciferase activity using GloMax Discover Microplate Reader (Promega).

### Ha-NiV neutralizing Antibody Assay

Ha-NiV particles (45 μl) were pre-incubated with 15 μl serially diluted sera for 15-30 minutes at room temperature in 96-well plates, 15 μl HEK293T cells (5×10^4^ cells per well) were used for infection for 18 hours at 37 °C. Cells were lysed in 7.5 μl Cell Lysis Buffer (Virongy) and 25 μl Luciferase Assay Buffer (Virongy) were added for luciferase assays using GloMax Discover Microplate Reader (Promega).

### PNRT neutralization assay

Neutralization activity of serum antibodies was analyzed by plaque reduction neutralization test (PRNT) as described previously with an exception that NiVb was used in this study ^40^. Briefly, AGM sera were heat inactivated at 56°C for 1 hour and were serially diluted 1:25-fold for ten dilutions in Dulbecco’s Phosphorous buffered saline (DPBS, Gibco). Each dilution was incubated at 37°C and 5% CO2 for 1 hour with an equal volume of 1000 PFU/ml of NiVb diluted in DMEM (Gibco) with 2% FBS (Gibco). A 1000 PFU/ml NiVb in DMEM and only DMEM with 2% FBS were positive and negative controls respectively. A 200 µl of each dilution of incubated serum along with controls were added to confluent monolayers of NR Vero E6 cells in triplicates and plates were incubated for 1 hour at 37°C and 5% CO2. Plates were gently rocked frequently to prevent drying of monolayers. An overlay of 2.5% Avicel RC-591 microcrystalline cellulose and Carboxymethyl Cellulose Sodium (DuPont Nutrition & Biosciences) was mixed in 1:1 ratio with 2X Modified Eagle Medium (Temin’s modification, Gibco) and 10% FBS (Gibco). Plates were incubated for 4 days at 37°C and 5% CO2. After incubation period, the monolayers were fixed with 10% neutral buffered Formalin and stained with 0.2% aqueous Gentian Violet (RICCA Chemicals) in 10% neutral buffered formalin for 30 minutes, followed by rinsing. Plates were dried and plaques were counted manually. The half maximal inhibitory concentrations (IC_50_) were calculated using GraphPad Prism 8.

### Quantification and statistical analysis

Infection and luciferase assays were performed in triplicate. Luciferase expression was quantified with luciferase assay and the mean was the average value of the three luciferase assay readings. Standard deviations (SD) were determined using Microsoft Excel. Antibody neutralization activity was plotted using GraphPad Prism 7 and the IC_50_ values were calculated using GraphPad Prism 7.

